# SeqsLab: an integrated platform for cohort-based annotation and interpretation of genetic variants on Spark

**DOI:** 10.1101/239962

**Authors:** Ming-Tai Chang, Yi-An Tung, Jen-Ming Chung, Hung-Fei Yao, Yun-Lung Li, Yin-Hung Lin, Yao-Ting Wang, Chien-Yu Chen, Chung-Tsai Su

**Affiliations:** Atgenomix, Taipei, Taiwan; Genome and Systems Biology Degree Program, National Taiwan University and Academia Sinica, Taipei, Taiwan; Graduate Institute of Medical Genomics and Proteomics, National Taiwan University College of Medicine, Taipei, Taiwan; Department of Bio-Industrial Mechatronics Engineering, National Taiwan University, Taipei, Taiwan; Education & Research Center for Bio-Industrial Automation, National Taiwan University, Taipei, Taiwan.

## Abstract

**Summary:** SeqsLab is a platform that helps researchers to easily annotate and interpret genetic variants derived from a large quantity of personal genomes. It provides an integrated interface to annotate the variants based on curated databases as well as *in silico* estimation on the effects of the variants. SeqsLab adopts the scalable cluster computing framework, Spark, and incorporates several customized algorithms to speed up the process of variant annotation and interpretation. The key features of SeqsLab include efficient annotation on large structural variations, diverse combinations of variant filters, easy incorporation with a vast amount of public databases, and scalable architecture of analyzing hundreds of human whole genomes simultaneously.

**Availability and Implementation:** SeqsLab is implemented with JAVA. The generated annotation will then be stored in Elasticsearch for real-time query and exploratory analysis. SeqsLab can be accessed by web browsers and is freely available at http://portal.seqslab.net/.

**Supplementary information:** Supplementary data are available at *Bioinformatics* online.

## 1. Introduction

The advance of high-throughput sequencing (HTS) technologies over recent years has increased our ability to decode a personal genome in a short time. Raw reads generated by HTS technologies are mapped back to the reference genome before calling different types of variants, including single nucleotide variations (SNV), small insertions or deletions (indels), and large structural variations (SVs). Numerous tools, such as ANNOVAR (Yang and Wang 2015), VEP (McLaren, Pritchard et al. 2010), SnpEff (Cingolani 2012), VAT (Habegger, Balasubramanian et al. 2012), and vcfanno (Pedersen, Layer et al. 2016), have been developed to annotate those acquired variants by comparing the called variants with comprehensive publicly available datasets. The huge amount of available data not only results in time-consuming processes but also makes researchers hard to extract useful information from the enormous output results. While most of the annotator tools focus on improving the annotating speed and flexibility, the tedious filtering and interpretation processes are left to the researchers as the downstream analysis without help from an integrated platform. Some web services such as Oncotator (Ramos, Lichtenstein et al. 2015) can do both annotation and visualization of the observed variants, but usually set limitation on the number of query variants or types of annotation databases in order to reduce computing resources and database curation efforts. Here, we present a genomic analysis platform (Fig. 1), SeqsLab, to tackle the abovementioned issues. SeqsLab provides lightning-fast annotation workflow, comprehensive structural variant annotation, user-friendly graphical interface, and plenty variant filtering utilities to facilitate personal genome annotation and interpretation. As we are facing a huge number of variants, a user-friendly filtering tool is essential when researchers are trying to explore the variants of interest.

**Fig. 1.**
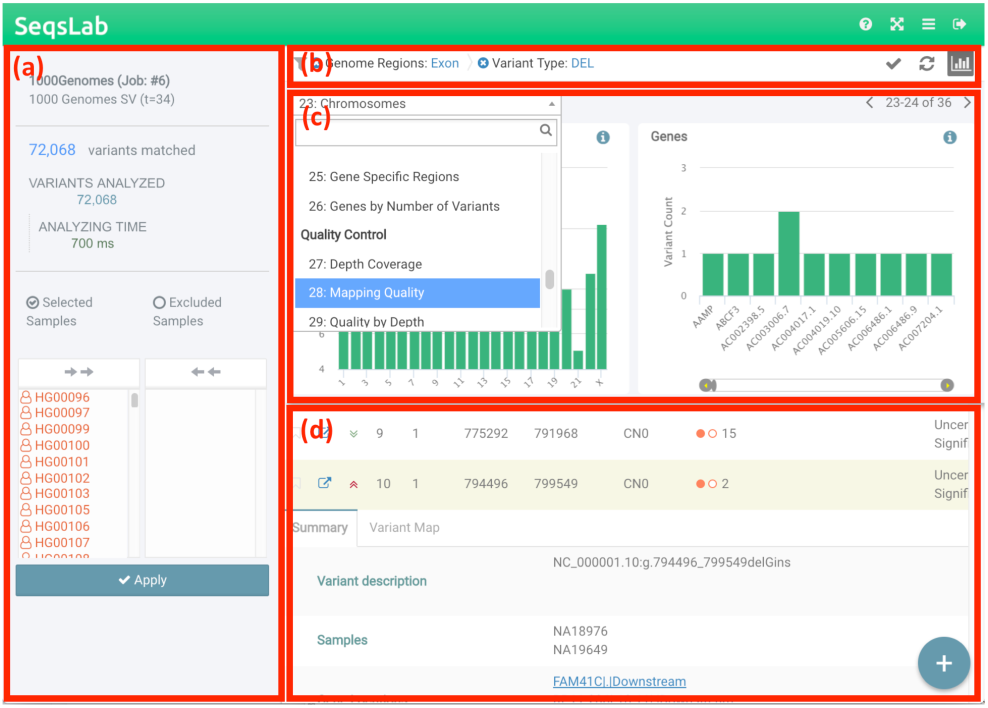
User interface of SeqsLab filters. **(a)** The summary page that reveals the current status of the annotation job, including the number of matched variants, the number of analyzed variants, analyzing time, and the IDs of the samples used. **(b)** The filters and the rules that have been applied. **(c)** The graphical representation of the filters in SeqsLab. The drop-down list contains all the available filters for the current job. **(d)** The list of brief descriptions for the matched variants.

## 2. Features and Methods

SeqsLab first leverages Spark in-memory computing framework (Gupta, Dutt et al. 2003) to annotate SNVs with all the curated databases. Then, indels and SVs are annotated by utilizing a proprietary graph algorithm to identify overlapped records against the curated databases containing known structural variations and genomic contexts. All the annotation results will be formed into JSON format and inserted into the Elasticsearch database (Divya and Goyal 2013). SeqsLab uses the Django framework to construct the web server and integrates jBrowse (Buels, Yao et al. 2016) as the genome viewer to show the locations of a selected variant and its relationships to the interacting partners on the genome. For whole genome sequencing (WGS) data of an individual, there will usually be 3~5 millions of variants identified after compared with the reference genome (hg19/hg38) using a standard pipeline. In order to quickly narrow down the suspects of the causal variants into a smaller set, we designed and implemented several filter plugins. Users can use different combinations of the well-designed filters to refine the results easily.

### 2.1 Curation of publicly available databases

The most updated versions of many popular databases for annotation were curated in SeqsLab. In Table 1, the curated databases were categorized into four groups (population, genomic context, clinical context, and functional context). Population databases majorly aim at collecting the sequencing data of normal individuals from various populations and locations in order to provide the allele frequency of each variant in different populations. Genomic context databases, such as GENCODE (Consortium 2004, Harrow, Frankish et al. 2012), ENCODE, and DENdb (Ashoor, Kleftogiannis et al. 2015), reveal the locations that have been already annotated as genes, transcripts, and regulatory elements. Clinical databases contain collections of ever-reported pathogenic variants on certain diseases. As Mendelian diseases are concerned, several Mendelian disease related databases such as deafness variant dataase (DVD) (Shearer, Eppsteiner et al. 2014), ClinVar (Landrum, Lee et al. 2014), and Leiden Open Variation Database (LOVD) (Fokkema, Taschner et al. 2011) were included. Furthermore, many well-known functional annotation databases (e.g. dbNSFP (Liu, Wu et al. 2016) and dbscSNV (Liu, Wu et al. 2016)) were included for general purposes. The original data downloaded from the public databases is kept in SeqsLab as well; so the users can always check the raw information from the public databases whenever needed. More details of the databases can be found in the supplementary file.

**Table 1.**
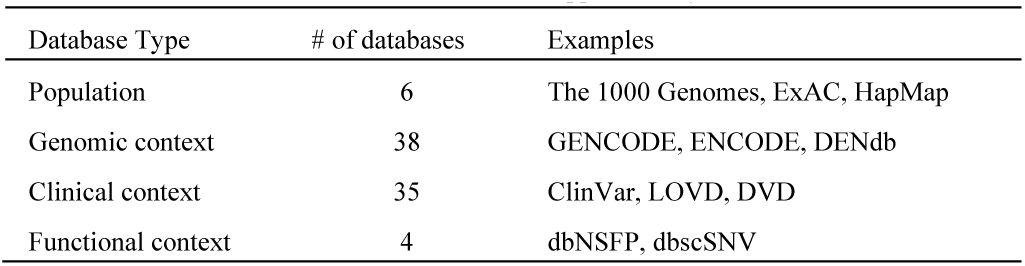
Four categories of the curated databases. The table contains the summary of the databases. More details can be found in the supplementary file.

### 2.2 System Flow

SeqsLab partitions the annotating procedures into four modules as illustrated in Fig. 2. First, the Data Validation module starts to verify all the selected VCF files, merges them by using the BCFtools package (Li, Handsaker et al. 2009) and then uploads the merged VCF file to HDFS. The remaining three modules are executed on Spark for parallel computation. All of point mutation variants are extracted from the merged VCF file and mapped to the curated databases for annotation in the Spark Join module. Then, indels and structural variations are extracted and submitted to the Spark Overlap Graph module, along with the variants that cannot be annotated in the Spark Join module. In the Spark Overlap Graph module, the variants are sorted together with the curated records of structural variations and genomic contexts by their chromosomes and positions. The details of this module will be described in the next section. All of the variants will be analyzed by the Variant Pathogenicity module, where the compound heterozygous variants can be identified if phasing information is provided. Last, all of variants will be stored and indexed in Elasticsearch (Gormley and Tong 2015), a distributed full-text search engine.

**Fig. 2.**
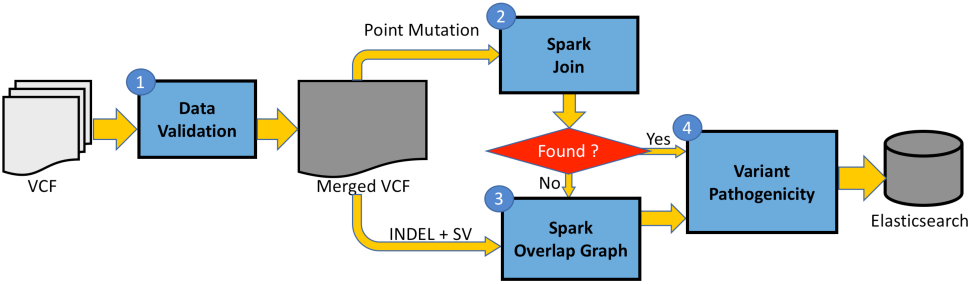
The system flowchart of SeqsLab.

### 2.3 Structural variant annotation algorithm

The differences between SNV annotation (the Spark Join module) and SV annotation (the Spark Overlap Graph module) is that SNV annotation usually seeks for an exact match of both the loci and alternative allele, while SV annotation should look for all the overlapping events and partners. All of records in the curated databases overlapped with the structural variations in the query VCF file must be comprehensively reported. One of the strategies used in previous SV annotators, such as ANNOVAR, is to seek for any overlaps with the query variants. This strategy is both time and memory consuming. Instead, SeqsLab utilizes a proprietary graph algorithm developed by Atgenomix (Atgenomix 2016) to speed up the process of SV annotation. The main concept of the algorithm is to sort both of the annotation data and SVs in the given VCF file by their positions, to construct overlapping graphs parallelly on Spark GraphX, and then to traverse all the records for the variants in order to generate all the relationships that should be labeled as relevant.

### 2.4 Variant filtering plugins

SeqsLab designed plenty of plugins to filter out the variants that are less potential to be the causal variants and to highlight the variants that have effects on protein functions or gene expression regulation. In Fig. 1(b) and Fig. 1(c), users can apply different filtering strategies based on their applications or research interests. The filtering interface is designed to be graphical and interactive. Therefore, users can apply any of the filtering plugins to remove unwanted variants interactively. SeqsLab provides seven types of filters, which are general property, medical relevance, variant effect, splicing event, genome position, quality control, and tissue specificity, to fulfill different aspects of filtering strategies. All the necessary data will be indexed during the annotation process. In this regard, the filtering process is relatively fast such that the users can explore as many filter combinations as possible in near real time.

## 3. Example usage

To demonstrate performance and scalability, SeqsLab uses two population-scale sequencing data. One is all structural variations identified in the 1000 Genomes Project (Via, Gignoux et al. 2010, Sudmant, Rausch et al. 2015); the other is a collection of whole genome sequencing data (30X coverage by Illumina HiSeq 2500) from Taiwan Biobank (contract number: TWBR10411-03).

### 3.1 Structural variation annotation

SeqsLab has the ability to provide comprehensive information for variant annotation and interpretation in an integrated platform. Since most existing tools do not support sufficient functionality for analyzing structural variations, here we demonstrated the power of SeqsLab on annotating structural variations for a large cohort efficiently and effectively. For this purpose, we downloaded the publicly available VCF files of structural variations from the 2,504 samples in the 1000 Genomes Project. The consensus set of the structural variations, which were called by three different SV callers, covers 68,818 loci, involving 72,068 variants in total. As an example, we adopted the filter combination: “Genome Regions=Exon”, “Variant Type=DEL” and “Novel Variants=True” along with “Inheritance=Autosomal Dominant” to identify the deletions potentially have functional effects to the proteins or sequence effects with respect to the reference genome. This filter combination quickly reduced the number of variants from 72,068 to 23, including several variations that might have functional effects on important genes.

### 3.2 Scalability of SeqsLab

We further performed scalability test on a set of whole genome sequencing data (30X coverage by Illumina HiSeq 2500) from Taiwan Biobank (contract number: TWBR10411-03). All of the sequencing data were processed by BWA (Li and Durbin 2009) and GATK UnifiedGenotyper (McKenna, Hanna et al. 2010). In average, each sample contains around four million of point mutations and seven hundred thousand of non-point mutations (including indels and SVs). In Fig. 3, the execution time of the four modules versus the number of samples is presented. As the trend of execution time over an increase of samples, the execution time of the Data Validation module (colored in blue) is roughly linear with the number of samples because of the I/O bound on merging VCF files. By leveraging in-memory computing of Spark, the execution time of the Spark Join module (colored in orange) and the Variant Pathogenicity module (colored in yellow) are under linear with the number of samples. Actually, their execution time are linearly related to the number of variants because of Spark join operation. By using Atgenomix’s proprietary graph algorithm on GraphX, the execution time of the Spark Overlap Graph module (colored in green) also performed linearly with the number of samples. According to Fig. 3, 128 WGS VCF files (around 600 GB) can be comprehensively annotated with more than 80 public available databases in less than 15 hours. Therefore, SeqsLab shows highly scalability on population-based annotation, especially for structural variations. For future enhancement on the Data Validation module, BCFtools can be trivially replaced by using the Spark framework to improve its performance.

**Fig. 3.**
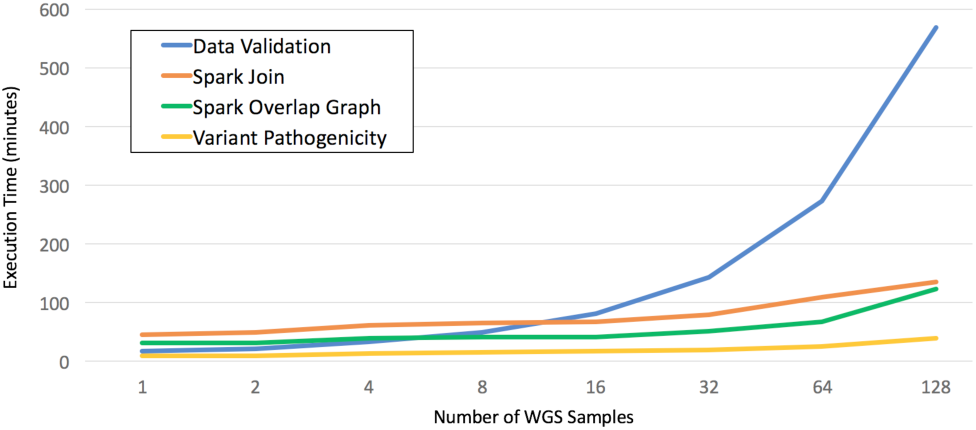
Performance Evaluation on SeqsLab. Execution time vs. Number of samples, running on a cluster of nine commodity servers (each node has four Intel Xeon quad-cores processors and is equipped with 64 GB memory)

## 4. CONCLUSIONS

SeqsLab is an integrated platform that leverages the Spark framework and uses proprietary graph algorithm to accelerate the annotation process of different types of variants, including SNVs, indels and structural variations. The rich information provided by the databases curated in SeqsLab can offer the users an efficient way to filter variants, to examine the distribution of the selected variants on each plugin, and to visualize the comprehensive information from curated databases. Because the sequencing cost drops continuously, a user-friendly platform that is capable of analyzing the whole genome data in a rapid and accurate manner will be highly appreciated. Though SeqsLab is currently focusing on variant annotation, some of the upstream analyses such as read mapping and variant calling will be integrated on the same platform in the near future.

## Acknowledgements

We thank Yun-Chian Cheng for critical reading of this article, and Chun-Pei Cheng for early construction of annotation.

none declared.

